# DUSP6 is upregulated in metastasis and influences migration and metabolism in pancreatic cancer cells

**DOI:** 10.1101/2024.12.20.629199

**Authors:** Mariana T. Ruckert, R. McKinnon Walsh, Bailey A. Bye, Austin E. Eades, Wei Yan, Filip Bednar, Jiaqi Shi, Costas A. Lyssiotis, Michael N. VanSaun, Vanessa S. Silveira

**Affiliations:** Department of Genetics, Ribeirao Preto Medical School, University of Sao Paulo, Ribeirao Preto, SP, 14040-901, Brazil; Department of Cancer Biology, University of Kansas Medical Center, Kansas City, KS, 66160, US; Department of Molecular & Integrative Physiology, University of Michigan, Ann Arbor, MI, 48109, US; Department of Surgery, University of Michigan, Ann Arbor, MI, 48109, US; Rogel Cancer Center, University of Michigan, Ann Arbor, MI, 48109, US; Department of Pathology & Clinical Labs, University of Michigan, Ann Arbor, MI, 48109, US; Department of Internal Medicine, Division of Gastroenterology and Hepatology, University of Michigan, Ann Arbor, MI, 48109, US; Massachusetts General Hospital Cancer Center, Krantz Family Center for Cancer Research, Boston, MA; Harvard Medical School, Boston, MA, 02114, US

## Abstract

Pancreatic ductal adenocarcinoma (PDAC) is a highly aggressive malignancy characterized by KRAS mutations in approximately 95% of cases. Despite recent advancements with KRAS inhibitors, therapeutic resistance has emerged, and combination approaches are needed. In particular, it is important to understand how downstream signaling of KRAS supports PDAC growth. For example, DUSP6, a dual-specificity phosphatase that modulates ERK1/2 phosphorylation and RAS pathway activity, has emerged as an important regulator of KRAS-MAPK signaling. Transcriptomic analyses demonstrate that DUSP6 is markedly overexpressed in PDAC tumors compared to non-tumoral pancreatic tissue. Single-cell RNA-seq reveals its upregulation in epithelial tumor cells, with further elevation in metastatic lesions relative to primary tumors. This upregulation correlates with the quasi-mesenchymal/squamous molecular subtype, and clinically, high DUSP6 expression is associated with poorer overall survival. Gene set enrichment analyses of metastatic samples indicate that DUSP6 is linked to pathways involved in cell migration and metabolism. To elucidate DUSP6’s functional roles, stable knockdown of DUSP6 in PDAC cell lines resulted in increased ERK/MAPK activation and altered migratory capacity. Metabolic profiling showed enhanced basal glycolysis following DUSP6 suppression. However, combined inhibition of glycolysis and DUSP6 downregulation did not affect the migratory phenotype, indicating that glycolytic alterations do not drive migration. These findings highlight the dual and independent roles of DUSP6 in modulating migratory capacity and glycolysis in PDAC. This study underscores the significance of DUSP6 as a potential therapeutic target and provides new insights into its contributions to PDAC progression.

**SIGNIFICANCE STATEMENT:** DUSP6 plays dual and independent roles in pancreatic cancer, regulating both migration and glycolysis. Its upregulation in metastasis is associated with poor prognosis and more aggressive phenotypes, highlighting its clinical relevance. Targeting DUSP6 represents a potential therapeutic strategy to disrupt KRAS-MAPK signaling and target key pathways driving PDAC progression.

## INTRODUCTION

Pancreatic ductal adenocarcinoma (PDAC) accounts for over 90% of pancreatic cancer cases and is one of the most lethal forms of invasive cancer globally, ranking as the third leading cause of cancer-related mortality [1–3]. Despite significant advancements, the current 5-year survival rate remains low, at 13% [3]. Late detection of PDAC primarily stems from vague symptoms, lack of early diagnostic markers, and resistance to chemotherapy [4, 5]. Complete resection remains the sole curative option, yet it is unattainable for the majority of patients; 80– 90% of patients present with locally advanced or metastatic disease upon diagnosis [6, 7]. Metastasis in PDAC commonly affects the liver, peritoneum, and lungs [8–12]. Studies indicate that not all circulating tumor cells successfully initiate secondary tumors, suggesting the requirement for genetic and/or epigenetic changes [13]. Mutations such as those occurring in *Kras* are prevalent in both primary and metastatic tumors, with certain passenger mutations showing a preference for metastatic sites [11, 12]. For example, loss of DPC4 (SMAD4) is more frequent in metastatic cases, correlating with increased invasiveness [14]. Gene expression alterations are also important considerations in this context; however, to date, most studies have only documented changes between non-neoplastic pancreatic and tumor tissues [15].

*Kras* mutations are found in approximately 95% of PDAC cases [16] and are the first genetic alterations that arise in precancerous lesions. These mutations alter protein conformation and ultimately lead to a prolonged activation state and, consequently, downstream activation of signaling pathways such as the ERK/MAPK pathway [17]. In tumor cells, ERK phosphorylation drives MYC activation through subsequent phosphorylation, leading to a myriad of effects such as uncontrolled cell proliferation, increased survival, and metabolic rewiring [18, 19]. Although it is intuitive to believe that KRAS constitutive activation is imperative for tumors, its hyperactivation induces oxygen reactive species (ROS) production, and cells are forced to upregulate their antioxidant systems to maintain ROS homeostasis. However, failure to buffer intracellular ROS accumulation can result in oncogene-induced senescence and cytotoxicity [20]. Therefore, fine-tuned KRAS-MAPK activation is required for tumor cells to survive and concurrently evade the detrimental effects of their metabolic adaptations.

DUSP6 is a dual-specificity phosphatase that targets serine and threonine residues, particularly dephosphorylating ERK1/2 and downregulating the ERK/MAPK pathway [21]. While DUSP6 is reported to act in a negative feedback loop to downregulate ERK/MAPK activation, it has also been reported that ERK1/2 phosphorylates DUSP6 to target this protein for subsequent proteasomal degradation [22]. Its role in cancer is context dependent, acting either as an oncogene or as a tumor suppressor [23]. DUSP6 was previously shown to be upregulated in pancreatic intraepithelial neoplastic (PanIN) lesions (early stage of PDAC tumorigenesis) and downregulated in invasive carcinoma, particularly in poorly differentiated subtypes [24]. This phenomenon has been reported to occur due to hypermethylation of *DUSP6* promoter, leading to abrogation of gene expression and increased activity of the ERK/MAPK pathway [25]. The complex coordination of the KRAS-ERK-DUSP6 axis and the timely regulation of DUSP6 expression throughout tumorigenesis prompted us to investigate the role of this molecule in PDAC metastasis.

In this study, we demonstrated that *in silico* gene expression and RNAscope studies in human samples revealed upregulation of *DUSP6* in metastatic samples compared to that in primary tumors and non-neoplastic pancreatic tissue. Gene set enrichment analysis (GSEA) of human metastatic PDAC samples revealed enrichment of pathways related to cell migration and metabolism. *In vitro* assessments demonstrated *DUSP6* downregulation in a KRAS mutant cell line, which correlates with increased ERK/MAPK activation, diminished proliferation, and migration capacity; however, this effect was absent in KRAS wild-type cells. Functionally, *DUSP6* downregulation led to elevated basal glycolysis, indicating its involvement in glucose metabolism. However, the combined inhibition of *DUSP6* and glycolysis did not affect the migratory phenotype of the KRAS mutant cell line. Our findings suggest that, while DUSP6 influences migration capacity and glucose metabolism, these effects may not be directly correlated in this context.

## MATERIALS & METHODS

### Cell lines and culture

MIA PaCa-2 (RRID:CVCL_0428), PANC-1 (RRID:CVCL_0480), AsPC-1 (RRID:CVCL_0152), BxPC-3 (RRID:CVCL_0186), Capan-2 (RRID:CVCL_0026), and SW1990 (RRID:CVCL_1723) were purchased from the ATCC (Manassas, VA, USA). K8484 (RRID:CVCL_XD12) and DT8082 (RRID:CVCL_XD11) were acquired from David Tuveson’s laboratory (Cold Spring Harbor Laboratory) [20]. P4313 was acquired from Andy Lowy (UC San Diego) [21]. PKT62 is a murine pancreatic tumor cell line derived from a PKT (*LSL-Kras*^G12D/+^, *TGFBR2*^fl/fl^, *Ptf1a*^Cre/+^) mouse [26] and kindly provided by Michael VanSaun (University of Kansas Medical Center). KPC CAFs and PKT CAFs cell lines were isolated from the respective mouse tumors and validated by western blotting. Briefly, tumors were collected, digested, minced and cells were sorted with EpCAM beads (BioLegend Inc., cat #741444). EpCAM^−^ cells were kept in culture and until full confluence, protein samples were isolated and fibroblastic phenotype was confirmed by the presence of α-SMA and absence of CK19 and Ras^G12D^. Capan-2, AsPC-1 and SW1990 were cultured in high glucose (4.5 g/L) RPMI-1640 medium (ThermoFisher Scientific, Waltham, MA USA cat #A1049101) supplemented with 10% heat-inactivated fetal bovine serum (FBS, Atlanta Biologicals/RD Systems, Atlanta, GA USA) and antibiotic-antimycotic (Thermo Fisher Scientific cat# 15240062). The K8484 cell line was cultured in high-glucose (4.5 g/L) Dulbecco’s Modified Eagle Media (Thermo Fisher Scientific, cat# 11-995-073) supplemented with 5% heat-inactivated FBS (R&D Systems-Atlanta Biologicals) and antibiotic-antimycotic (Thermo Fisher Scientific). All other cell lines were maintained in high-glucose DMEM supplemented with 10% FBS and antibiotic-antimycotic. Cells were maintained in culture conditions (37°C, 5% CO_2_ and humid atmosphere) as recommended by the ATCC, utilized within 15 passages from purchase and regularly tested for mycoplasma.

### Plasmids and gene knockdown

To generate *DUSP6* knockdown cells, short hairpin RNA (shRNA) constructs targeting three independent regions of DUSP6 mRNA (#1 5’-CTGTGGTGTCTTGGTACATTG-3’; #2 5’-TCTAATCCAAAGGGTATATTT-3,’ #3 5’-ATTCGGCATCAAGTACATCTT-3’) were obtained from Millipore (Burlington, MA, USA). The empty vector pLKO.1 puro (RRID:Addgene_8453) was used as a control. Packaged lentiviruses were generated by transfecting HEK293T (RRID:CVCL_0063) cells with the respective plasmids and the Trans-Lentiviral™ Packaging Mix (Open Biosystems) to produce viral supernatants, which were subsequently concentrated with Lenti-X™ concentrator (TakaraBio, cat #631231). Next, 2.5×10^5^ pancreatic cancer cells were seeded in 6-well plates and incubated for 24h with lentiviral suspension. Successfully transduced cells were selected with puromycin treatment (1-2.5µg/mL) until complete cell death was achieved in the non-transduced cells.

### RNA isolation and RT-qPCR

Briefly, 10^6^ cells were seeded in 6-well plates and incubated overnight. The next day, cells were washed with 1X PBS and harvested using QIAzol Lysis Reagent (Qiagen, GmbH, cat #79306). Samples were stored at −80°C until total RNA was isolated using the Direct-zol RNA Miniprep kit (Zymo Research Corporation), according to the manufacturer’s protocols. RNA quantification and quality analyses were performed using NanoDrop 2000 (Thermo Fisher Scientific Inc.). For cDNA, 1.5-2µg of RNA was processed using the High-Capacity cDNA Reverse Transcription Kit (Applied Biosystems, cat #4368814) according to the manufacturer’s instructions. The product was further diluted 1:10 before storage at −20°C. Real-time PCR analysis was carried out with the 2X SYBR Green qPCR Master Mix (ApexBio Technology, cat #K1070) on a ViiA 7 Real-Time PCR System (Applied Biosystems) using the following primers: human *DUSP6* (Fwd: ATGGTAGTCCGCTGTCCAAC/Rev: ACGTCCAAGTTGGTGGAGTC), mouse *Dusp6* (Fwd: TGTTTGAGAATGCGGGCGAGTT / Rev: ACAGTTTTTGCCTCGGGCTTCA), human *ACTB* (Fwd: TCGTGATGGACTCCGGTGAC/Rev: CGTGGTGGTGAAGCTGTAG), and mouse *Actb* (Fwd: GTGACGTTGACATCCGTAAAGA / Rev: GCCGGACTCATCGTACTCC). Each sample was run in triplicate using the 2^−ΔΔCT^ method (Livak & Schmittgen, 2001) and the fold-change was evaluated relative to normal samples and determined using *ACTB/Actb* levels as a reference.

### Western blotting

Briefly, 10^6^ cells were plated in 6-well plates, incubated overnight, and collected the next day for protein isolation. Cells were washed with 1X PBS, lysed in ice-cold RIPA buffer (1X RIPA [Cell Signaling Technology]; 10mM NaF; 1mM PMSF), sonicated, and incubated on ice for 10 min. The cells were then centrifuged at 16,000 × *g* for 10 min at 4°C. Protein quantification was performed using the Pierce™ BCA Protein Assay Kit (Thermo Fisher Scientific), following the manufacturer’s instructions. For western blotting assays, 10 µg of total protein was loaded in each lane of a 10% polyacrylamide gel and subjected to electrophoresis. Proteins were transferred to nitrocellulose membranes using the semi-dry method on a Trans-Blot Turbo Transfer System (Bio-Rad). Specific proteins were detected using the following primary antibodies: P-ERK1/2 (Cell Signaling Technology, cat #4370, RRID:AB_2315112, [link](https://scicrunch.org/resolver/AB_2315112)), ERK1/2 (Cell Signaling Technology, cat #4695, RRID:AB_390779, [link](https://scicrunch.org/resolver/AB_390779)), E-cadherin (Cell Signaling Technology, cat #3195, RRID:AB_2291471, [link](https://scicrunch.org/resolver/AB_2291471)), N-cadherin (Cell Signaling Technology, cat #13116, RRID:AB_2687616, [link](https://scicrunch.org/resolver/AB_2687616)), and β-actin (Cell Signaling Technology, cat #12262, RRID:AB_2566811, [link](https://scicrunch.org/resolver/AB_2566811)) and MKP-3 (Santa Cruz Biotechnology, cat #sc-137246, RRID:AB_2095347, [link](https://scicrunch.org/resolver/AB_2095347)). Secondary antibodies used were Peroxidase AffiniPure F(ab’)₂ Fragment Donkey Anti-Mouse IgG (H+L) (cat #715-036-151, RRID:AB_2340774, [link](https://scicrunch.org/resolver/AB_2340774)) and Donkey Anti-Rabbit IgG (H+L) (cat #711-036-152, RRID:AB_2340590, [link](https://scicrunch.org/resolver/AB_2340590)) from Jackson ImmunoResearch Laboratories Inc. Images were obtained using FluorChem M (Bio-Techne) or a Sapphire FL Biomolecular Imager (Azure), and analyzed using ImageJ software (NIH, Bethesda, MD, USA).

### Immunohistochemistry staining

Healthy pancreata, primary tumor, and metastatic samples were collected from wild-type C57BL/6J (RRID:IMSR_JAX:000664) mice, *LSL-Kras*^G12D/+^, *Ptf1a*^Cre/+^ (KC), *LSL-Kras*^G12D/+^, Tp53^R172H/+^, *Ptf1a*^Cre/+^ mouse pancreatic tissue (KPC) maintained in a C57BL/6J background. Male and female mice were used for experiments and mice ages varied from 5.9 to 86.6 weeks. Samples were collected, embedded in paraffin, and sectioned into 4μm sections. The paraffin sections were processed as follows: 15 min in xylene (2x); 5 min in 100% ethanol (2x) and 5 min in 75% ethanol. Finally, the sections were washed 3x in PBS at room temperature (RT). Antigen retrieval was performed by boiling the sections in 10 mmol/L sodium citrate solution for 10 min and waiting for the sections to cool down and achieve RT. Next, the sections were washed 3x with PBS, quenched with 3% H_2_O_2_ for 15 min, and blocked with 5% BSA for 20 min. Whole-slide scanning was performed using a PANNORAMIC Scan II scanner (3DHISTECH). Immunohistochemical staining was performed by iHisto Inc. (Salem MA; iHisto.io) using an anti-DUSP6 antibody (Abcam, cat #ab76310, RRID:AB_1523517, link). Ten fields per independent section from each condition were randomly selected for quantification using the QuPath software [27]. The study weas performed under protocols 2005N000148 and 2019N000116 approved by the IACUC at MGH, and at Pharmaron Inc. and Champions Oncology under approved protocols.

### RNA *in situ* hybridization

RNA in situ hybridization was performed using the RNAscope® technology (Advanced Cell Diagnostics, Inc.). Briefly, tumors were collected from the mice, sectioned, and immediately fixed in 10% buffered formalin for 24h. Tumor sections were preserved in 70% ethanol, embedded in paraffin blocks, cut into 5 µm sections, and placed in pre-treated slides. The sections were freshly cut for the assay and dried at RT to preserve RNA integrity. Sections were baked for 1h at 60°C and then deparaffinized using Histo-Clear (Electron Microscopy Science, cat #6411004) and 100% ethanol. Next, sections were treated with RNAscope® Hydrogen Peroxide for 10 min at RT, followed by a target retrieval step with RNAscope® 1X Target Retrieval Reagent for 15 min, and finally, RNAscope® Protease Plus treatment for 30 min at 40°C. Finally, we used the RNAscope® 2.5 HD Duplex Reagent Kit, following the manufacturer’s protocol. Specific probes used in this assay were Mm-Dusp6 (cat #429321-C1), Mm-Krt19-C2 (cat #402941-C2) and Mm-Pdgfrb-C2 (cat #411381-C2). The sections were imaged at 20x magnification using an EVOS™ M5000 Imaging System (Thermo Fisher Scientific).

### Clinical samples

All the human patient samples described in this study were collected from patients undergoing pancreas resection due to pancreatitis, cystic neoplasms or PDAC from 2002 to 2015 at the University of Michigan Health System. Collection of these samples was approved by the Institutional Review Board at the University of Michigan (IRB number: HUM00098128).

### RNA *in situ* hybridization combined with immunofluorescence multiplex staining

Tissue microarray (TMA) sections were obtained, set up and processed as described in [28]. Fluorometric in-situ hybridization was performed according to the manufacturer’s (Advanced Cell Diagnostics, Inc.) instructions. Briefly, FFPE preserved TMAs were cut in 5 µm sections, and placed in pre-treated slides. Sections were freshly cut for the assay and dried at RT to preserve RNA integrity. Sections were baked for 1h at 60°C, deparaffinized in xylene three times for 15 mins, washed in 100% ethanol twice for 3 mins and air dried. Slides were treated with pretreatment 1, 2 and 3 buffers, rinsed in double-distilled water (ddH_2_O) and incubated with the DUSP6 (cat #1032061-C1) or control probes for 2h at 40°C in a humidity chamber. Slides were then treated with Amp solutions 1-6 before the fluorogenic substrate Opal 690 TSA (Akoya Biosciences, Inc.) was applied to the slide. After a 10 min incubation, the slides were washed and then subjected to dual immunofluorescent staining on a Ventana Discovery Ultra stainer (Roche Diagnostics, North America) with smooth muscle actin (BioCare Medical, cat #001, RRID:AB_2910606, [link](https://scicrunch.org/resolver/AB_2910606), 1:100, 60 mins, Opal 570) and pan-cytokeratin (Abcam, cat #ab9377, RRID:AB_307222, [link](https://scicrunch.org/resolver/AB_307222), 1:200, 20 mins, Opal 520). Slides were mounted using Prolong Gold with DAPI and scanned using a Vectra Polaris™ (Akoya Biosciences, Inc.) whole slide scanner. Images were analyzed using the de-array TMA tool in QuPath and DAPI was used to identify individual cells in each core.

### Proliferation assay

Cell proliferation was assessed by ethynyldeoxyuridine (EdU) incorporation. Briefly, 2.5-5×10^4^ cells were seeded in 24-well plates in complete media and incubated overnight. The next day, the media was replaced with the appropriate treatments and incubated for 18 h. At this point, EdU was added to the wells at a final concentration of 10µM and cells were incubated for an additional 6h. At the end of this period, the cells were trypsinized, harvested, and fixed with 10% buffered formalin overnight at 4°C with gentle agitation. The next day, the cells were washed with PEB-T (1x PBS, 2mM EDTA, 1% BSA, 0.5% Triton-X) and subsequently with PB (1x PBS, 1% BSA). Next, the cells were resuspended in Click-It Reaction (H_2_O, 100mM Tris pH 8.5, 1mM CuSO_4_, 1µM azide-dye) and 100mM ascorbic acid. Cells were incubated with 0.5µg/mL propidium iodide (PI), washed twice, and resuspended in PEB (1x PBS, 2mM EDTA, 1% BSA) for flow cytometry analysis. The cells were analyzed for 530 nm and 695 nm emission using an Attune NxT Flow Cytometer (Thermo Fisher Scientific). FCS files were analyzed with FlowJo™ v10.8 Software (BD Biosciences) and the percentage of EdU-positive cells was taken from the PI-positive single cells.

### Migration assay

First, 2-4×10^5^ cells were seeded into 24-well plates and cultured until they reached full confluence. Then, a scratch was made at the center of the plate using a 200µL pipette tip. Wells were washed twice with 1X PBS to discard all debris, and media with appropriate treatments were added. The plates were photographed at 0 and 24h using an EVOS™ M5000 Imaging System (Thermo Fisher Scientific). Scratch area was calculated using the Wound Healing Coherency Tool (Montpellier Ressources Imagerie) on ImageJ (NIH, Bethesda, MD, RRID:SCR_003070, [link](https://scicrunch.org/resolver/SCR_003070)) and results are expressed as healing percentage – the difference in scratch area between time 0 and time 24.

### In silico analysis

*DUSP6* expression analysis was performed using publicly available datasets obtained from The Cancer Genome Atlas (TGCA_PAAD), Genotype-Tissue Expression (GTEx, normal pancreatic tissue) and Gene Expression Omnibus (GEO, RRID:SCR_005012, [link](https://scicrunch.org/resolver/SCR_005012); GSE62452 [29], GSE28735 [30], GSE15471 [31], GSE71729 [32] and GSE224564 [33]). Overall survival analysis was performed by Gene Expression Profiling Interactive Analysis (GEPIA, RRID:SCR_018294, [link](https://scicrunch.org/resolver/SCR_018294)) using the UCSC Xena project (http://xena.ucsc.edu) datasets. Differential expression of *DUSP6* in the epithelial and stromal compartments was performed using GSE93326 [34] with the R2 Genomics Analysis and Visualization Platform. The latter was also utilized for accession of Bailey *et al.* [35] dataset since ICGC Data Portal Repository was discontinued. Finally, *DUSP6* expression was correlated with Metastatic Potential (MetMap 500 [36]) at the DepMap Portal using the Cancer Cell Line Encyclopedia (CCLE). We specifically filtered the cell lines for Exocrine Adenocarcinoma, which corresponds to PDAC cell lines. Gene set enrichment analysis was conducted in a cohort of metastatic samples from the publicly available dataset GSE71729 [32] using the hallmark gene set collection (version 2024) available at GSEA 4.3.3. software (RRID:SCR_003199, [link](https://scicrunch.org/resolver/SCR_003199)) [37]. Single-cell analysis was performed using a previously published [38] pancreatic tissue single-cell atlas ([link](https://pascadimaglianolab.shinyapps.io/SC_Pancreas_Atlas/)).

### Glycolysis stress test

Glycolysis rates were determined using the Seahorse XF Glycolysis Stress Test Kit (Agilent, Santa Clara, CA, USA), following the manufacturer’s protocol. Briefly, 2-4×10^4^ cells were seeded into a 96-well plate and incubated overnight. The next day, cells were washed twice with Seahorse XF DMEM (pH 7.4 (2mM glutamine) and a final volume of 180µL of the same was added to each well. The extracellular acidification rate (ECAR) was measured (mpH/min) at 11 time points with sequential injections of 10mM Glucose, 1µM Oligomycin and 100mM 2-Deoxy-D-glucose (Thermo Scientific, cat. #AC111980010). For nuclear staining, 5µg/mL Hoechst was added to the oligomycin solution. The ECAR values were calculated and normalized to the cell count number using an automated imaging and cell counting workflow. For normalization, the ECAR value per 10^3^ cells was set, and the Agilent Seahorse XF system automatically calculated and generated normalized results.

### Lactate assay

For lactate measurement, 2.5×10^4^ cells were seeded in 96-well plates and allowed to sit overnight in the incubator. The next day, the complete media in the plate was replaced with new media containing 0.5% FBS, and the cells were incubated for 24h. At the end of the experiment, culture plates were centrifuged at 450 × *g* for 5 minutes and 10µL of media was collected from each well for the subsequent assay. For lactate quantification, we utilized the Glycolysis Cell-Based Assay Kit (Cayman Chemical Company, Ann Arbor, MI, USA, cat #600450), following the manufacturer’s protocol. The absorbance was read at 490 nm using a Synergy H1 Multimode microplate reader (Bio-Tek/Agilent Technologies).

### Statistical analysis

Sample sizes were chosen to achieve a minimum of triplicate samples for all the experiments. For the assessment of statistical significance between two groups, Welch’s t-test was used for parametric data, while Mann-Whitne test was used for non-parametric data. For comparison across multiple groups, Brown-Forsythe and Welch ANOVA test or mixed-effects analysis with Šídák’s multiple comparisons test were used when appropriate for parametric data; for non-parametric data, Kruskal-Wallis test were performed. Data normality was determined based on Q-Q plots distribution. All statistical tests were performed using GraphPad Prism 9 software (RRID:SCR_002798, [link](https://scicrunch.org/resolver/SCR_002798)), and the results were considered significant at α = 0.05. * P < 0.05, ** P <0.01, *** P < 0.001, and **** P < 0.0001.

## RESULTS

### *DUSP6* overexpression is correlated with advanced stage PDAC

To investigate the role of DUSP6 in metastasis, we first confirmed its expression in tumor samples. To evaluate *DUSP6* expression in PDAC samples, we used three distinct in silico datasets available on public platforms (GSE62452 [29], GSE28735 [30] and TCGA/GTEx). Our analysis revealed that *DUSP6* was upregulated in primary tumor samples compared to non-tumoral pancreatic tissue across all analyzed datasets (Figure 1A-C). Based on these findings, we aimed to assess *DUSP6* expression in metastatic cases. Utilizing the *in silico* dataset GSE71729 [32], we compared *DUSP6* expression across a cohort comprising 134 non-tumor tissue samples, 145 primary tumor samples, and 62 metastatic samples derived from various secondary sites. Notably, we observed that *DUSP6* was significantly overexpressed in metastatic samples compared to both primary tumor samples (P < 0.0001) and non-tumorous pancreatic tissue (Figure 1D). Considering that *DUSP6* is overexpressed in primary tumors and metastatic samples, we hypothesized that there is a possible correlation between DUSP6 expression and patient prognosis. To answer this question, we first assessed its expression in the PDAC molecular subtypes. We performed *in silico* analysis utilizing PDAC molecular subtyping [35, 39]. We took advantage of the datasets from Badea *et al.* (GSE15471) and Bailey *et al.* [35] and analyzed *DUSP6* expression in distinct PDAC subtypes. In the Badea *et al.* cohort (Figure 1E), we observed that *DUSP6* was overexpressed in the quasi-mesenchymal subtype compared to non-tumoral pancreatic tissue (P < 0.0001) and the exocrine-like subtype (P = 0.0211). Likewise, in the Bailey *et al.* cohort (Figure 1G), we observed DUSP6 overexpression in the squamous subtype compared to that in the pancreatic progenitor (P = 0.0065) and immunogenic (P = 0.0197) subtypes. Next, we performed the same analysis using another subtyping paradigm [32]; however, no differences were observed (Figure 1F). The same subtyping was used to stratify samples in the GSE224564 [33] (Figure 1H) dataset, and we observed overexpression of *DUSP6* in the basal-like subtype compared to the classical subtype (P = 0.0292). The quasi-mesenchymal, squamous, and basal-like subtypes share similar molecular signatures and worse prognoses than the other PDAC subtypes. Collectively, these results suggest that DUSP6 is enriched in this tumor phenotype. To establish a link between DUSP6 expression and ERK1/2 activation, we performed a correlation analysis between *DUSP6* expression and *MAPK1* expression in quasi-mesenchymal samples from GSE15471 [31]. We observed a moderate positive correlation in this context (R = 0.6787, P = 0.02), suggesting that DUSP6 overexpression may be required to counteract MAPK signaling (Figure S1A).

**Figure 1.**
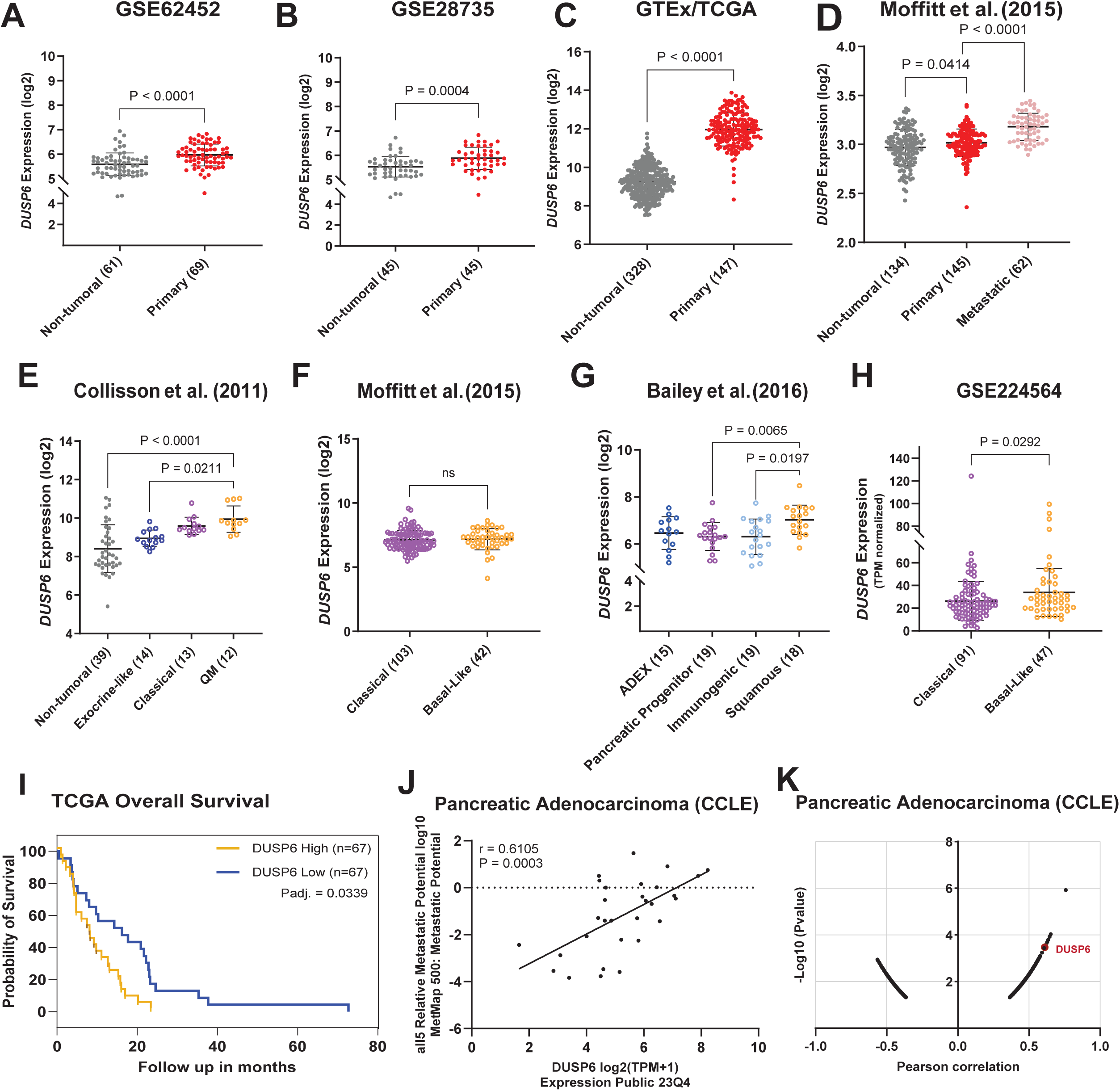
*DUSP6* overexpression is correlated to tumor metastatic potential and worse prognosis. *In silico* datasets from GSE62452 [29] **(A)**, GSE28735 [30] **(B)** and GTex/TCGA **(C)** assessing *DUSP6*’s differential expression between non-tumoral pancreatic tissue and primary tumor tissue; **(D)** GSE71729 [32] assessing *DUSP6* expression between non-tumoral pancreatic tissue, primary and metastatic tumor tissue; *In silico* gene expression analysis utilizing **(E)** Collisson [39], **(F)** Moffitt [32], **(G)** Bailey [35] and **(H)** GSE224564 [33] datasets to assess *DUSP6* expression among PDAC molecular subtypes; **(I)** Kaplan-Meier (Log-rank) plot comparing *DUSP6* expression and patients’ overall survival using the TCGA dataset. Padj. = 0.0339, N = 67 for each group; **(J & K)** Pearson correlation analysis between *DUSP6* expression in CCLE PDAC cell lines and their metastatic potential (MetMap 500 [36]). R = 0.6105, P = 0.0003. ADEX: aberrantly differentiated endocrine– exocrine; QM: quasi-mesenchymal. (A, H) Mann-Whitney unpaired test. (B, C, F) Welch’s unpaired t-test. (D, G) Brown-Forsythe and Welch unpaired ANOVA test. (E) Kruskal-Wallis unpaired test. * P < 0.05; ** P < 0.01; *** P < 0.001; **** P < 0.0001.

Supporting the latter, our analysis of TCGA dataset revealed that high *DUSP6* expression in PDAC patients correlated with worse overall survival (P = 0.0058; Figure 1I). Despite the counterintuitive nature of *DUSP6* overexpression in metastatic samples, given the prevailing dogma suggesting its negative regulation of the ERK/MAPK pathway, our findings were substantiated by further correlation analysis conducted with 30 PDAC cell lines from the Cancer Cell Line Encyclopedia (CCLE) and the Metastasis Map (MetMap 500 [36]). This analysis revealed a moderate positive correlation between *DUSP6* expression and metastatic potential in these cell lines (R = 0.611; P = 0.0003; Figure 1J). Furthermore, this correlation analysis indicated that *DUSP6* gene expression ranked as the eighth strongest positive correlation for metastatic potential within this PDAC cell line cohort among a list of 1000 genes (Figure 1K). Collectively, these observations suggest that *DUSP6* overexpression in PDAC samples is associated with the poorest prognostic outcomes and correlates with progression to metastasis.

To validate these findings, we assessed DUSP6 expression in lesions across different stages of the disease using healthy murine pancreatic tissue, primary tumors, and liver metastasis samples. We observed a trend of temporal control of DUSP6 expression with disease progression, although the results were not statistically significant (Figure 2A). DUSP6 expression is upregulated during the acinar-to-ductal metaplasia (ADM) process and subsequently reduced in pancreatic intraepithelial neoplastic (PanIN) lesions. DUSP6 was again increased in primary tumor lesions and, finally, in liver metastasis samples. These results suggest that temporal control of DUSP6 expression is differentially utilized during PDAC tumorigenesis and progression, perhaps to fine-tune MAPK signaling.

**Figure 2.**
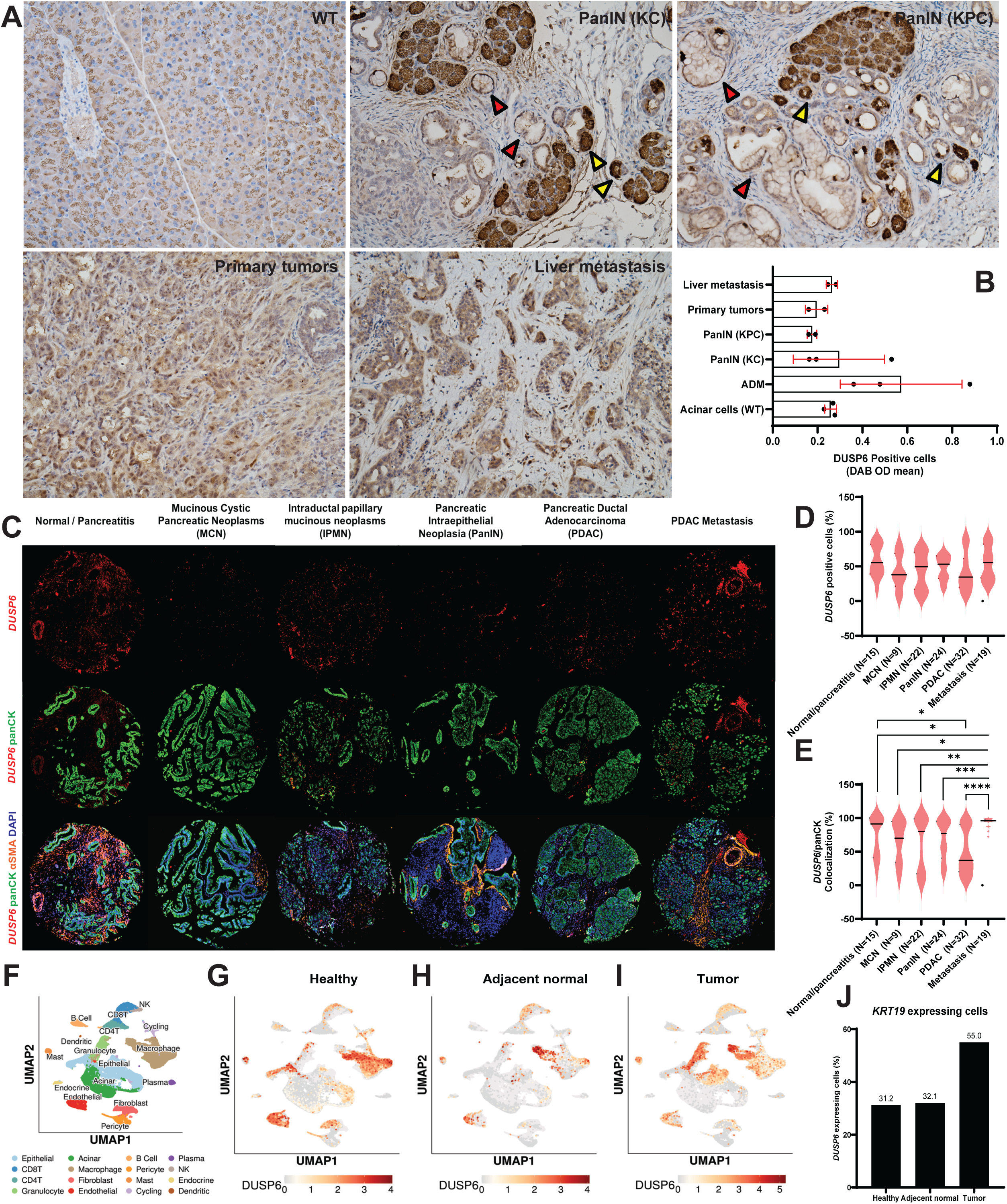
*DUSP6* is upregulated in PDAC epithelial cells in human and mouse tissue. **(A)** DUSP6 immunohistochemistry staining (20x magnification) in wild type (WT) pancreatic tissue (N=3), KC (N=3) and KPC (N=6) mice. Yellow arrows indicate acinar-to-ductal metaplasia (ADM) lesions and red arrows indicate Pancreatic Intraepithelial Neoplasia (PanIN) lesions. Quantification (B) shows DAB OD mean ± SD in DUSP6 positive cells from 10 randomly selected areas in each slide stratified by lesion grade; **C)** Multiplex *in situ* hybridization with *DUSP6* (red, RNAscope), panCK (green), αSMA (orange) and DAPI (blue). Images are representative of normal pancreatic tissue/pancreatitis, Intraductal papillary mucinous neoplasms (IPMN), Mucinous Cystic Pancreatic Neoplasms (MCN), Pancreatic Intraepithelial Neoplasia (PanIN), Pancreatic Ductal Adenocarcinoma (PDAC) and PDAC metastasis samples composed from 2 human TMA sections; **D)** Quantification of the percentage *DUSP6* positive cells normalized by total number of cells in each TMA core; **E)** Quantification of the percentage of cells that are positive for both *DUSP6* and panCK, normalized by total number of cells in each TMA core. Welch’s unpaired t-test; **F)** UMAP plot of all cells captured from single-cell RNA sequencing of eleven human tumor samples [28]. Cell types are identified by color. Following UMAPs show DUSP6 expression among cell types in healthy pancreatic tissue **(G)**, adjacent normal pancreatic tissue **(H)** and PDAC samples **(I)**; **J)** Percentage of cells co-expressing *DUSP6* and *KRT19* (epithelial marker) in healthy pancreatic tissue, adjacent normal pancreatic tissue and PDAC samples assessed by single-cell RNA-seq [28]. * P < 0.05; ** P < 0.01; *** P < 0.001; **** P < 0.0001.

PDAC has a highly complex microenvironment and is mainly composed of dense fibroinflammatory stroma consisting predominantly of activated stromal fibroblasts and immunosuppressive immune cell types. Our previous analyses were primarily performed on bulk tumors, where gene expression represents a composite of myriad cell types in the tumor microenvironment. Accordingly, we assessed whether DUSP6 overexpression was localized to specific cell types within the tumor. To answer this question, we examined *DUSP6* expression in different tumor compartments using GSE93326 [34]. Indeed, our analysis revealed that *DUSP6* was significantly upregulated in tumor cells compared to stromal cells (P < 0.0001; Figure S1B). We validated this analysis by assessing DUSP6 protein levels in four mouse PDAC cell lines and two mouse cancer-associated fibroblast (CAF) cell lines. The mouse cell lines exhibited varying levels of the protein, with DT8082 and K8484 demonstrating the highest DUSP6 levels, P4313 showing intermediate levels, and PKT62 displaying the lowest levels (Figure S1C). Subsequently, we investigated *Dusp6* expression in mouse tumor tissue using RNA in situ hybridization (RNA-ISH) technology. We examined *Dusp6* mRNA levels in tumor sections obtained from *LSL-Kras*^G12D/+^, *Ptf1a*^Cre/+^ (KC), *LSL-Kras*^G12D/+^, Tp53^R172H/+^, *Ptf1a*^Cre/+^ (KPC) mice, co-stained for either *Krt19* (a tumor epithelial cell marker) or *Pdgfrb* (a stromal marker). As depicted in Figure 1SC, *Dusp6* (green dots) exhibited strong co-expression with *Krt19* (red dots, upper images), but not with *Pdgfrb* (red dots, lower images), in both genetic backgrounds. Consistent with the western blot analysis of protein levels, both KC and KPC pancreata demonstrated *Dusp6* overexpression in malignant epithelial cells, whereas stromal compartment expression remained low (Figure S1D). To further explore the link between DUSP6 expression and MAPK signaling, we leverage GSE93326 [34] to perform a correlation analysis between *DUSP6* and *MAPK1* in the epithelial and stromal compartments. Interestingly, we observed a moderate positive correlation in gene expression levels in the epithelial compartment (R = 0.6105, P < 0.0001) but not in the stromal compartment (P = 0.0321, P = 0.789) in this cohort (Figure S1E-F). Altogether, these data indicate that DUSP6 is upregulated in PDAC epithelial cells in the context of hyperactivated MAPK signaling.

To confirm whether these observations could be translated into human disease, we used a panel of human samples from two TMAs containing normal pancreatic tissue/pancreatitis, intraductal papillary mucinous neoplasms (IPMN), mucinous cystic pancreatic neoplasms (MCN), PanIN, PDAC, and PDAC metastasis. We differentially stained tissue compartments with duplex RNA-ISH specific markers using panCK for epithelial cells and αSMA for CAFs (Figure 2C). A general assessment of *DUSP6* positive cells throughout the cores revealed a trend similar to that observed in mouse tissue. Although not statistically significant, we observed a slight decrease in expression in PDAC primary tumors compared to that in normal pancreatic tissue and a subsequent increase in metastatic samples (Figure 2D). Furthermore, we observed a significant increase in *DUSP6* expression in epithelial cells throughout tumor progression, and once again, a notable decrease in *DUSP6* expression in PDAC primary tumors, followed by upregulation in metastatic samples (Figure 2E). These findings suggest that *DUSP6* timely upregulation is required for metastatic tumor cells in the foreign environment.

The observation that *DUSP6* expression is significantly lower in PDAC primary tumor cells is counterintuitive, considering what we previously observed in bulk RNA-seq datasets. Considering these previous observations and the complexity of the PDAC microenvironment, we decided to dig further into a publicly available pancreatic tissue single-cell atlas [38] to characterize *DUSP6* expression patterns in different cell types within the healthy donor pancreas, adjacent normal pancreatic tissue, and primary tumor microenvironment. Figure 2F depicts the UMAP plot of all the cell types identified in the bulk of human patient samples. Interestingly, we observed that regardless of the context, DUSP6 was majorly expressed in myeloid cells, with increased expression in macrophages (Figures 2G-I, Figure S2A). Nevertheless, when we look specifically at the epithelial cells, in these three contexts, we observe a 2.4-fold increase in the percentage of *DUSP6* expressing epithelial cells in PDAC compared to healthy pancreata (Figure S2A). Lastly, we performed co-expression analysis of *DUSP6* and *KRT19* and confirmed a 1.8 × increase in *DUSP6* positive cells in PDAC *KRT19* positive cells compared to healthy pancreata (Figure 2J). These results support *DUSP6* upregulation in epithelial cells within the tumor microenvironment. Interestingly, *in silico* analysis utilizing a bulk RNAseq dataset from Bojmar *et al.* [40] reveals that *DUSP6* is upregulated in liver biopsy samples from patients who eventually developed PDAC liver metastasis in comparison to those who did not in a 3-year follow up study (Figure S2B). Our results also indicate a role for DUSP6 in the immune cell compartment, which may explain the variations in DUSP6 expression observed in different PDAC human samples when evaluated through bulk RNA sequencing. Additionally, DUSP6 expression may influence the establishment of the pre-metastatic niche.

### *DUSP6* expression in metastatic samples correlates with decreased oxidative phosphorylation and increased glycolysis

To gain further insight into the functional significance of *DUSP6* overexpression in metastatic samples, we aimed to elucidate the pathways involved. To accomplish this, we revisited the GSE71729 [32] dataset and conducted gene set enrichment analysis using 62 metastatic samples. Intriguingly, our analyses revealed enrichment of several pathways associated with cell motility, epithelial-to-mesenchymal transition, and metabolism. Notably, these enrichments revealed a positive correlation between *DUSP6* expression and cell migration (Figure 3B), suggesting an upregulation of this process, along with a negative correlation between its expression and oxidative phosphorylation (Figure 3A), suggesting a downregulation of this pathway. Supporting these observations, Panther Pathway enrichment analyses conducted across two independent datasets (TCGA_PAAD and CPTAC_PDAC) highlighted Glycolysis (P00024) as one of the pathways most strongly correlated with *DUSP6* expression in PDAC across all three datasets (Table S1). Moreover, other enrichment analyses utilizing the CPTAC Protein dataset indicated a correlation between DUSP6 protein levels and the Pentose phosphate pathway (P02762), Glucose 6-phosphate metabolic process (GO:0051156), and Carbon metabolism (hsa01200) (Table S1). These findings prompted us to hypothesize that *DUSP6* overexpression may play a pivotal role in metabolic reprogramming of tumor cells during metastasis.

**Figure 3.**
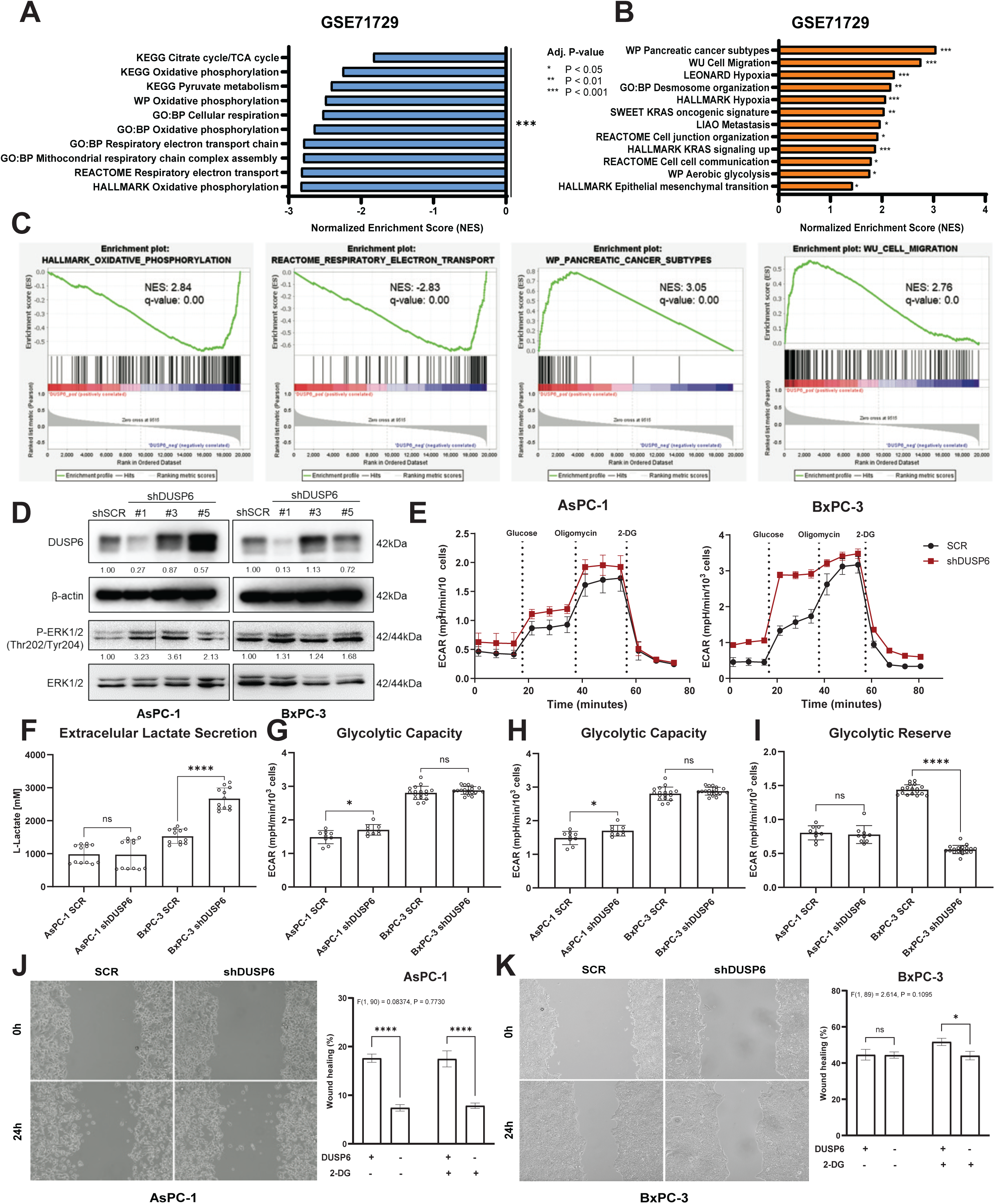
DUSP6 downregulation increases glycolysis in PDAC cell lines, but glycolysis inhibition does not rescue migratory phenotype. GSEA analysis of GSE71729 dataset showing pathways significantly negatively **(A & C)** and positively **(B & C)** correlated to *DUSP6* expression in metastatic samples; **D)** Western blot analysis of DUSP6 expression and ERK1/2 phosphorylation upon DUSP6 knockdown. Quantification shows DUSP6 expression normalized to β-actin, and P-ERK1/2 normalized to total ERK1/2 levels; **E)** Representative curves of extracellular acidification rates (ECAR) measured in AsPC-1 and BxPC-3 with sequential injections of glucose (10 mM), oligomycin (1 µM) and 2-DG (100 mM), respectively in a Seahorse assay; **F)** Basal extracellular lactate secretion levels upon DUSP6 knockdown in complete media; **G)** Basal glycolysis corresponds to glycolytic levels under basal conditions, no stimulation; **H)** Glycolytic capacity is measured after inhibition of ATP synthase (complex V of the mitochondrial respiratory chain); **I)** Glycolytic reserve corresponds to the difference between glycolytic capacity and basal glycolysis. Assessment of migratory capacity upon 2-DG treatment in AsPC-1 **(J)** and BxPC-3 **(K)** upon DUSP6 knockdown. Seahorse Glycolysis Stress Test was performed in 9-12 replicates in 2 independent experiments. Results represent mean ± SD of one representative experiment. L-Lactate assay was performed in 2 independent experiments with 6 replicates each. Results represent mean ± SEM. Welch’s unpaired t-test; Migration graphs show 3 independent experiments with 4-6 replicates each. Results represent mean ± SEM. Mixed-effects analysis with Šídák’s multiple comparisons test. *ns*: not significant; * P < 0.05; ** P < 0.01; *** P < 0.001; **** P < 0.0001.

### *DUSP6* downregulation reduces proliferation and migration in PDAC cell lines

Considering *DUSP6* upregulation in metastatic samples, we performed *DUSP6* knockdown in PDAC cell lines exhibiting the highest levels of the protein, one of which was derived from metastatic ascites (AsPC-1, Figure S3A). *DUSP6* knockdown was confirmed through gene expression and protein level analyses (Figure 3D, S3). As expected, *DUSP6* knockdown led to increased activation of ERK1/2, as confirmed by enhanced phosphorylation on western blotting (Figure 3D). Subsequently, considering that ERK1/2 activation elicits a proliferative phenotype, we evaluated cell proliferation using ethynyldeoxyuridine (EdU) incorporation assays under conditions of full serum (5%) and serum deprivation, designed to either stimulate or inhibit proliferation. Surprisingly, both AsPC-1 and BxPC-3 cell lines exhibited impaired proliferation after *DUSP6* knockdown. Notably, AsPC-1 cells displayed a more pronounced decrease, particularly under serum-starved conditions (P < 0.0001; Figure S4A). While BxPC-3 cells also showed significant proliferation impairment (P < 0.0001), they were less affected by serum starvation compared to AsPC-1 cells (Figure S4B).

To confirm our prior observations in human samples, we investigated the impact of *DUSP6* knockdown on cell migratory capacity. *In vitro* migration assays conducted within 24h of scratching (time 0) revealed distinct phenotypes among the cell lines. AsPC-1 cells exhibited a significant reduction in migratory capacity upon DUSP6 knockdown (P < 0.0001; Figure S4C), whereas BxPC-3 cells showed no significant effect (Figure S4D).

In summary, our findings suggest that DUSP6 plays a role in regulating both proliferation and migration capacity of PDAC cells, albeit with potential context-dependent differences.

### *DUSP6* downregulation increases glycolytic flux in PDAC cell lines

To ascertain DUSP6’s involvement in tumor cell metabolism, we first evaluated basal lactate secretion in extracellular media following *DUSP6* knockdown using a colorimetric assay. In this context, distinct phenotypes were observed in both cell lines: BxPC-3 cells exhibited a significant increase in lactate secretion compared to the control (P < 0.0001), whereas AsPC-1 cells showed no significant difference compared to the control (Figure 3F).

Considering that glucose is the primary source of energy for tumor cells *in vitro*, we sought to delineate how DUSP6 affects glucose metabolism in each cell line and conducted a metabolic assay to measure glycolysis activity in the absence of *DUSP6* (Figure 3E). Upon *DUSP6* knockdown and glucose injection, both AsPC-1 and BxPC-3 cells demonstrated a statistically significant increase in basal glycolysis (Figure 3G), consistent with observations from the lactate secretion assay. Following treatment with oligomycin, an inhibitor of ATP synthase (complex V), which shuts down mitochondrial respiration and makes cells dependent on glycolysis for ATP production, we observed a significant increase in glycolytic capacity in AsPC-1 cells upon *DUSP6* knockdown compared to the control (P< 0.05; Figure 3H), with no significant differences in glycolytic reserve (Figure 3I). This indicates that the absence of *DUSP6* alone leading to pronounced ERK1/2 activation pushes these cells towards their maximum glycolytic potential. Upon the addition of 2-DG, glycolysis was completely abolished under all conditions. These results demonstrate that *DUSP6* inhibition affects glycolysis in PDAC under basal metabolic conditions, but subsequent rescue experiments are required to determine the observed changes in glycolytic reserve.

### Glycolysis inhibition does not provide an additional impact cell migratory capacity in *DUSP6* depleted cells

To gain deeper insights into how metabolic changes induced by *DUSP6* knockdown affect cell behavior, we conducted a cell proliferation assay. By reducing fetal bovine serum in the media to minimize its metabolic influence, we assessed cell proliferation with and without 2-DG addition. Consistent with our previous findings, both AsPC-1 and BxPC-3 cells exhibited a significant reduction in proliferation upon *DUSP6* knockdown compared to their respective controls. Following 2-DG treatment, BxPC-3 cells displayed a notable decrease in proliferation under both conditions, with the reduction being more pronounced in the control cells (P < 0.0001; Figure S5B). In contrast, AsPC-1 showed no discernible effect on proliferation compared to the same conditions without 2-DG (Figure S5A).

Subsequently, to establish a link between the metabolic phenotype and metastatic potential of these cells, we conducted a migratory capacity assay combined with 2-DG treatment. *DUSP6* knockdown significantly reduced the migratory capacity of AsPC-1 cells, whereas no change was observed in BxPC-3 cells (Figure S4). Unexpectedly, treatment with 2-DG did not significantly alter the migratory phenotype observed in AsPC-1 cells, but significantly reduced BxPC-3 migratory capacity upon DUSP6 knockdown (Figure 3J-K). Interestingly, we noted that in the presence of an external source of pyruvate in the media, the cells maintained their glycolytic potential, even when treated with electron transport chain (ETC) inhibitors (Rotenone and Antimycin A) and 2-DG (Figure S5C). This suggests that pyruvate availability plays a critical role in the decision to sustain aerobic glycolysis or to feed the TCA cycle. These findings suggest that, under nutrient-rich conditions, both AsPC-1 and BxPC-3 utilize different energy sources for migration.

## DISCUSSION

DUSP6 is differentially expressed during PDAC tumorigenesis and progression [24]. However, little is known about the role of phosphatases in the development and establishment of metastasis. Our group observed that *DUSP6* was overexpressed in primary tumor samples compared to non-tumoral pancreatic tissues by analyzing *in silico* datasets and spatial transcript profiling. To further investigate the complex feedback loops involving DUSP6 and the ERK/MAPK signaling pathway, we evaluated *in silico* data to assess *DUSP6* expression in late-stage PDAC samples. We observed that *DUSP6* was overexpressed in metastatic samples compared to primary tumor samples by leveraging *in silico,* immunohistochemistry, and multiplex RNAscope analysis. Translational relevance for *DUSP6* expression was observed in survival data; TCGA analysis showed that high DUSP6 expression correlates with lower overall survival in PDAC patients. Finally, when we analyzed *DUSP6* expression among different PDAC molecular subtypes [35, 39], *DUSP6* overexpression was pronounced in the quasi-mesenchymal/squamous subtype, which is associated with a worse prognosis [41]. These data support the hypothesis that *DUSP6* overexpression confers a more aggressive phenotype, further impacting metastasis and worsening overall survival in PDAC.

Spatial localization is critical for investigating the functionality. Through *in silico* analysis of pancreatic tumor samples, we determined that *DUSP6* expression was upregulated in the epithelial compartment of the tumor rather than in the stromal compartment. The same pattern was observed in mouse models assessing protein presence by immunohistochemistry and RNA *in situ* hybridization in mouse pancreatic tumors. In human samples, we observed a significant decrease in *DUSP6* expression in PDAC epithelial cells versus normal pancreatic tissue, but increased expression in metastatic epithelial cells. By leveraging a single-cell RNA-seq Human Pancreatic Cancer Atlas, we were able to confirm that *DUSP6* expression is increased in PDAC epithelial cells. However, the results also highlighted macrophages as a subset of cells with important changes in *DUSP6* expression throughout tumorigenesis. Kidger *et al.* recently reported similar RNA *in situ* hybridization results using KC mice, demonstrating that *DUSP6* expression is found in ductal cells in PanINs, but not in stromal cells [42]. Our results confirmed that PDAC tumor cells overexpress *DUSP6*, suggesting the need for further investigation of its role in tumor development.

Previous studies on multiple tumor types have linked high DUSP6 expression to worse prognosis and metastasis [43–45]. *DUSP6* has alternatively been described as a tumor suppressor gene in PDAC. Furukawa *et al.* reported that downregulation of *DUSP6* coordinated PDAC progression via hypermethylation of its promoter [24, 25]. Kidger *et al.* also proposed a tumor suppressor role for *Dusp6*, observing that KCD6^−/−^ animals develop a higher number of poorly differentiated tumors and liver metastases [42]. In their model, *Dusp6* was constitutively absent, leading to a permanent state of Erk1/2 activation, which would predictably result in more aggressive tumors and more metastatic foci. Our current hypothesis is that *DUSP6* suppression might allow tumor initiation, while its expression is required for metastatic cells to re-establish in a new environment. Reinforcing this idea, *DUSP6* overexpression appears to correlate with the more invasive quasi-mesenchymal subtype.

To investigate this, we subjected PDAC cell lines to stable *DUSP6* knockdown and evaluated their proliferation and migration capacities. Surprisingly, we did not find consensus regarding the observed phenotypes. All statistically significant changes in proliferation indicated an impairment in this capacity. This observation is counterintuitive because it is widely known as a dogma in the field that ERK1/2 activation leads to increased cell proliferation. However, it is also known that tumor cells require fine-tuning of ERK1/2 activation, in which too much MAPK signaling can bypass the fitness advantage and become detrimental to cell growth [46]. Similarly, Gutierrez-Prat *et al.* recently reported that depletion of DUSP4, another ERK1/2 negative regulator, reduces proliferation by driving “oncogene overdose” in BRAF- and NRAS-mutant melanoma cell lines [47]. The migration phenotype varied among the cell lines, complicating any definitive conclusions on this topic. We hypothesized that this variability might be due to natural compensatory mechanisms within the cells, resulting from dependence on this signaling pathway. AsPC-1 cells carry two alleles of the *KRAS*^G12D^ mutation, which is the most commonly found among PDAC patients and, therefore, more clinically relevant. BxPC-3 cells present with wild-type *KRAS* which might explain the differences observed between these two cell lines. A recent study by Bulle *et al.* showed that upon MAPK pharmacological inhibition, *DUSP6* is one of the top downregulated genes, demonstrating that *DUSP6* expression correlates with ERK/MAPK activation. Supporting this evidence, we observed that *DUSP6* was significantly upregulated in tumor cell lines harboring a *KRAS* mutation compared to tumor cells driven by other mutations in the CCLE dataset (Figure S6). Bulle *et al.* also demonstrated that upon MAPK inhibition, DUSP6 was targeted for proteasomal degradation, leading to an increase in HER2 activation [48]. Interestingly, this is the first time that DUSP6 has been described as a negative regulator of HER2, since it was previously thought to be ERK-specific. This discovery shifts the paradigm of DUSP6’s functional role and raises the possibility that it may be involved in the negative feedback for other signaling pathways. Additionally, it is important to consider that DUSP6 is not the only phosphatase acting as a regulator of ERK phosphorylation. Therefore, its downregulation may elicit compensatory responses from other molecules, ultimately offsetting its absence. Ito and colleagues recently demonstrated a co-dependence of DUSP4 and DUSP6 in MAPK-pathway-driven tumors as a mechanism of drug resistance [49]. In this regard, we highlight that many DUSPs, among other proteins, are involved in ERK1/2 negative feedback control and have been shown to act synergistically to prevent oncogene-induced senescence [23]. Similarly, we found a positive correlation between DUSP6 and MAPK1 expression in tumor cells, but not in stromal cells, in a human cohort, suggesting that these cells are also dependent on DUSP6 for ERK1/2 modulation; we cannot, however, rule out the involvement of other DUSPs in the process. Moreover, the BxPC-3 cell line does not harbor a *KRAS* mutation, and thus, its tumorigenic behavior does not rely on this condition. The observed phenotype in this cell line might be driven by mechanisms different from those initially hypothesized. Conversely, the AsPC-1 cell line, a metastatic PDAC line harboring a *KRAS* mutation, more closely mimics the genetic background that we aimed to investigate. In this cell line, we observed a significant impairment in migration capacity upon *DUSP6* knockdown and increased ERK1/2 activation.

To understand the mechanisms by which *DUSP6* affects the metastatic potential of PDAC cells, we performed gene set enrichment analysis using the metastatic cohort of samples from the GSE71729 [32] dataset. This analysis indicated a positive correlation with cell migration and a consistent negative correlation with oxidative phosphorylation. Consequently, we conducted a glycolysis stress test and observed that the absence of *DUSP6* led to increased glycolytic function in PDAC cell lines. This upregulation of glycolysis was confirmed by the increased extracellular acidification rate upon glucose supplementation (in the absence of pyruvate), which could be completely reversed by the addition of 2-DG. This shift towards a more glycolytic phenotype was expected based on previous literature showing that *KRAS*^G12D^ increases glucose uptake and glycolytic flux through MAPK activation in PDAC cells: Ying and colleagues demonstrated that increased glycolytic flux induces the non-oxidative pentose phosphate pathway (PPP), which is involved in processes such as nucleic acid synthesis and is closely related to cell proliferative capacity [18]. However, in this context, we observed that although glycolysis was induced upon *DUSP6* knockdown, the cells showed a decreased proliferative capacity, a phenotype that is consistent among the cell lines. This behavior might be linked to inherent differences in cell metabolism among PDAC cell lines, as it has been previously reported that cells can present distinct metabolic profiles despite their growth rates [50]. Interestingly, Hsu *et al.* have demonstrated that DUSP6 is required for T-cell receptor-mediated glycolysis, and that in the absence of DUSP6, T cells show higher dependence on glutamine and fatty acids to fuel the TCA cycle [51]. These findings reinforce DUSP6’s involvement in metabolic reprogramming but highlight the context-dependency of the observed phenotypes.

Finally, we hypothesized that the induction of glycolysis through *DUSP6* inhibition would decrease the migratory potential of these cells, which could be rescued by the addition of 2-DG, leading to irreversible glycolysis inhibition. We treated the cells with 2-DG and evaluated their migratory capacity but did not observe significant changes with the combination of *DUSP6* and glycolysis inhibition. Previous studies have demonstrated that the induction of glycolysis leads to increased metastatic potential due to the promotion of epithelial-to-mesenchymal transition (EMT) [52, 53]. In the present study, we demonstrated that while cells acquire a more glycolytic phenotype upon *DUSP6* inhibition, their migratory capacity is abrogated. However, inhibition of glucose flux failed to rescue the observed phenotype. We hypothesized that, in a context where pyruvate is available in the culture media, cells can bypass 2-DG-mediated glycolysis inhibition. To estimate the effect of glycolysis induction on the metastatic behavior of *DUSP6* knockdown cells, it is recommended to closely control nutrient availability in the media. While it is tempting to extrapolate our observations to the decreased metastatic potential of the cells, we must reiterate that metastasis encompasses a variety of complex processes involving both intrinsic and extrinsic signals [54]. To further understand DUSP6’s role in metastatic potential, it will be necessary to explore other mechanisms involved in metastasis, such as invasive potential and the ability of these cells to adapt to a new environment, which is highly correlated with their metabolic reprogramming capacity.

## CONCLUSIONS

This study provides new insights into the role of DUSP6 in PDAC metastasis and metabolism. We demonstrated that DUSP6 inhibition elicits cell-specific effects on migratory potential, while inducing glycolysis in PDAC cells. However, we were unable to establish a direct correlation between the two processes. Future studies should include *in vivo* metastatic assays upon DUSP6 inhibition, as well as metabolomic analysis, to better understand how PDAC cells reprogram their metabolism in the absence of DUSP6 to thrive and survive in a secondary niche.

## ACKNOWLEDGEMENTS

We would like to acknowledge Dr. David Tuveson (CRUK) for supplying the murine K8484 cell line used in these studies: Dr. Andrew Lowy (UCSD) for supplying the P4313 cell line; Dr. Dafydd Thomas, at the Research Histology and Immunohistochemistry Core (University of Michigan), for conducting the RNA-ISH multiplex assay; the Merchant Lab (University of Miami) for use of the PKT62 cell line; and the Flow Cytometry Core Laboratory at KUMC, which is sponsored, in part, by the NIH/NIGMS COBRE grant P30 GM103326 and the NIH/NCI Cancer Center Grant P30 CA168524.

## AUTHOR’S CONTRIBUTIONS

MTR: methodology, data collection, formal analysis, visualization, paper writing, and editing; RMW: data collection, formal analysis, and editing; BBB, AEE, WY, FB, JS: data collection and analysis; CAL: supervision, methodology, data analysis, editing; MNV: supervision, methodology, data analysis, paper writing, and editing; VSS: funding acquisition, project administration, conceptualization, supervision, methodology, data analysis, paper writing, and editing.

## COMPLIANCE WITH ETHICAL STANDARDS

All experiments involving mice were approved by the Ethics Committee on Animal Research at Ribeirao Preto Medical School, University of São Paulo (protocol# 016/2018), and were conducted in accordance with the Guidelines of the Brazilian College of Animal Experimentation.

