## Supplementary Figures S1-6 for "DUSP6 is upregulated in metastasis and influences migration and metabolism in pancreatic cancer cells"

### Supplementary figure 1

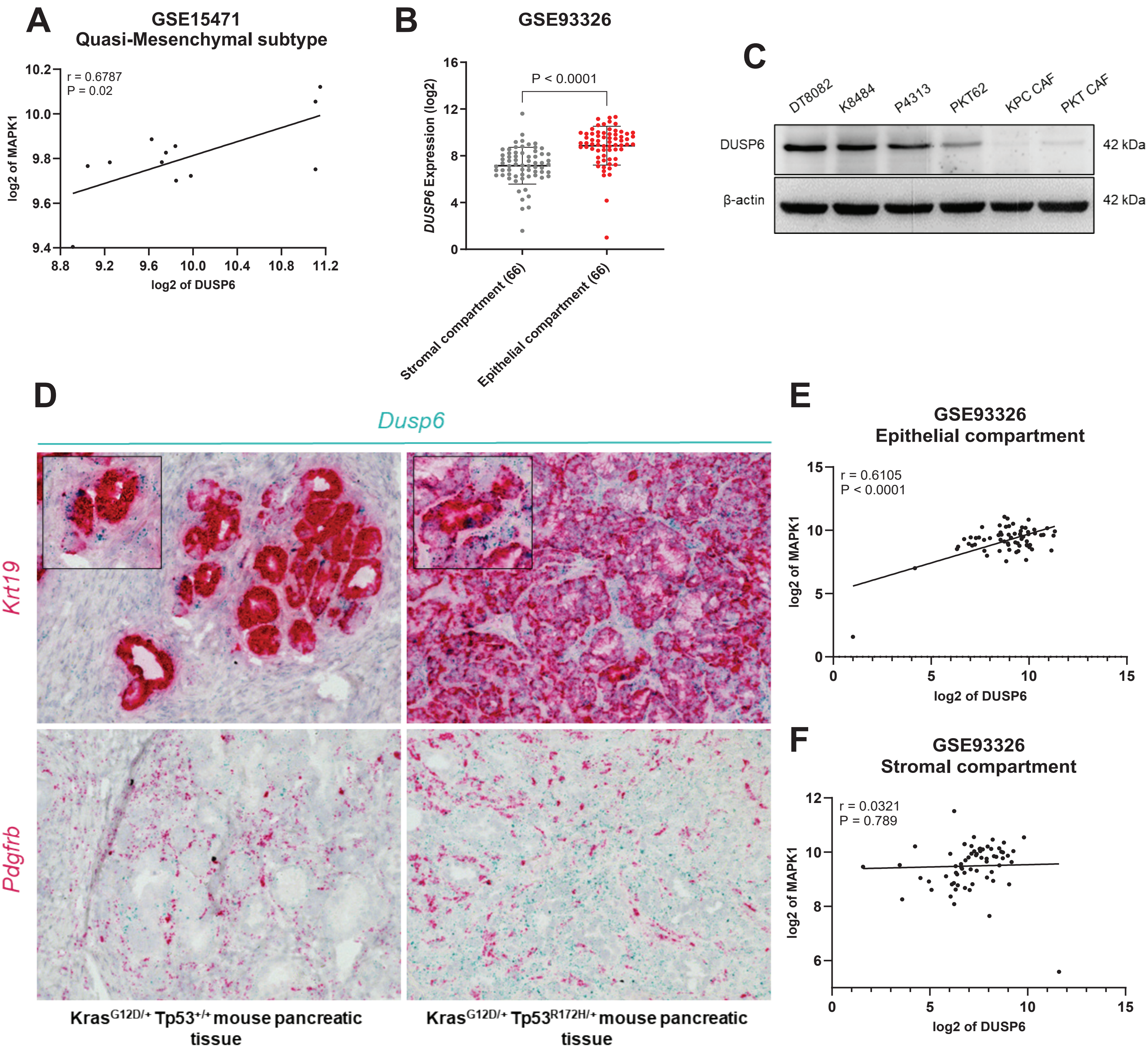

### Supplementary figure 2

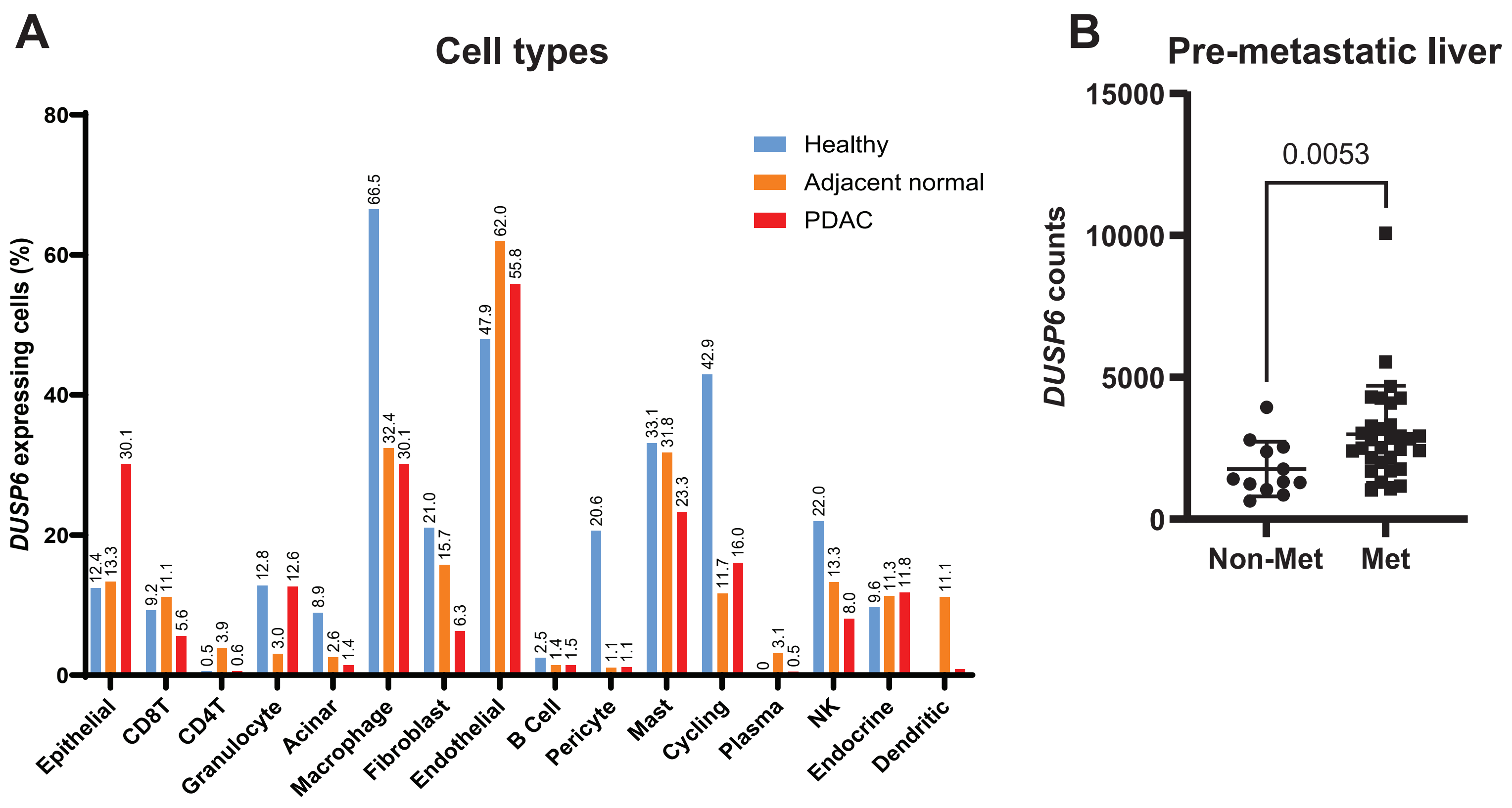


### Supplementary figure 3

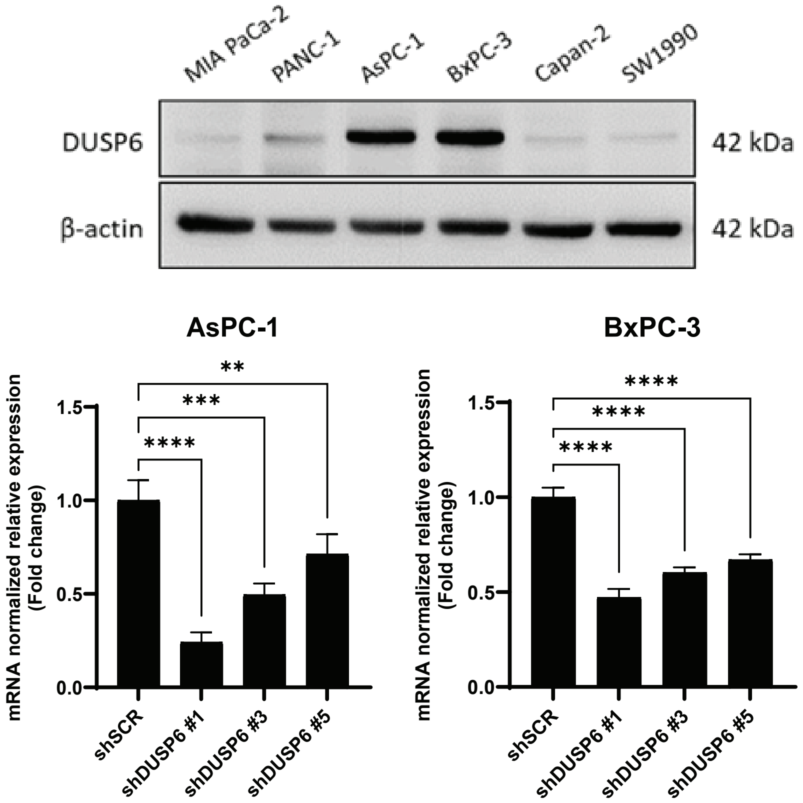

### Supplementary figure 4

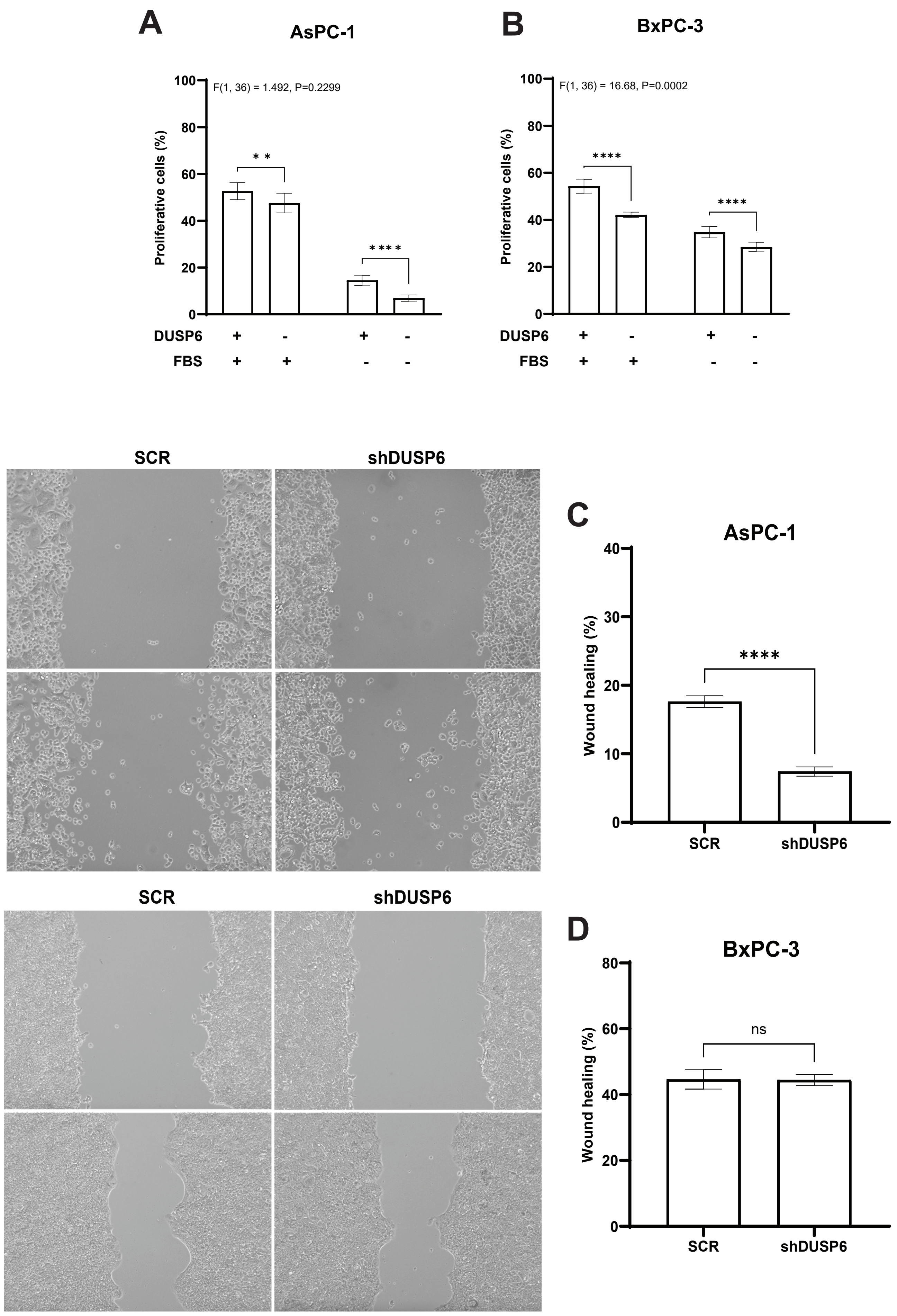

### Supplementary figure 5

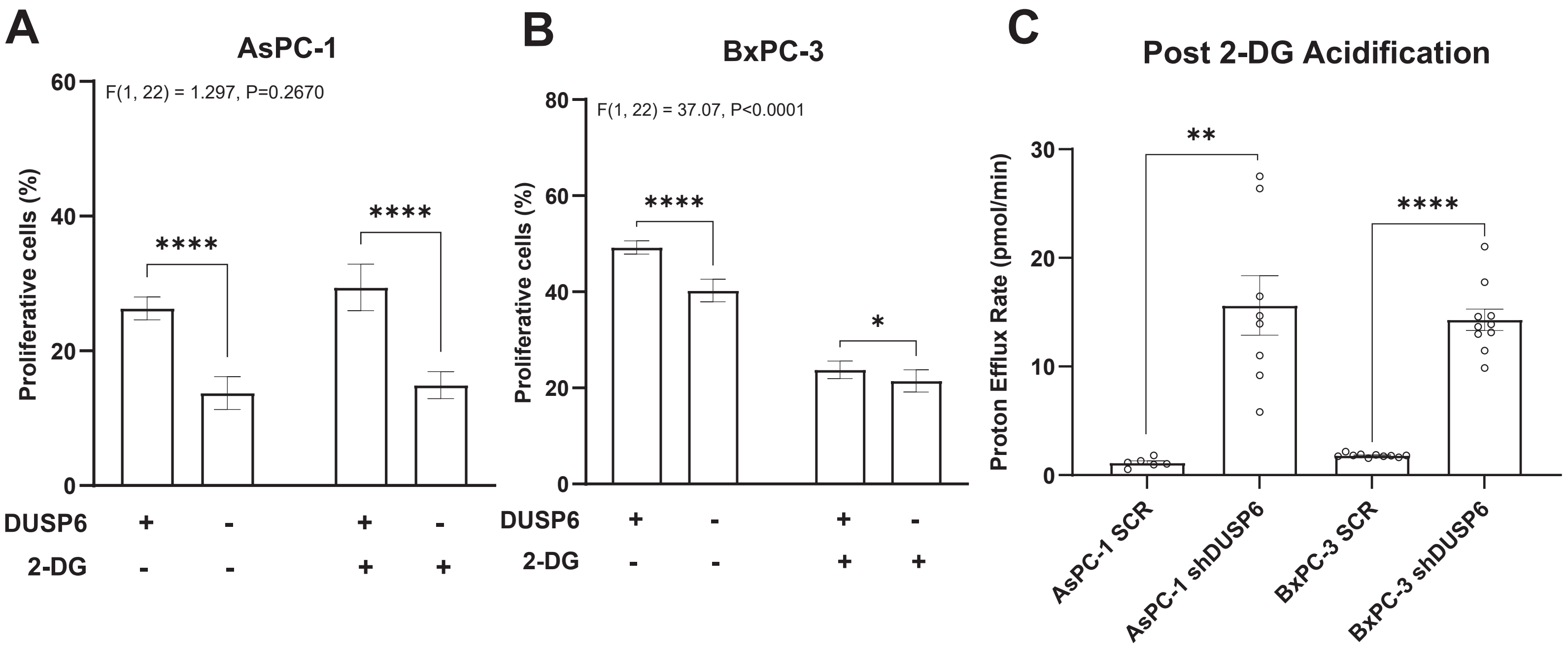


### Supplementary figure 6

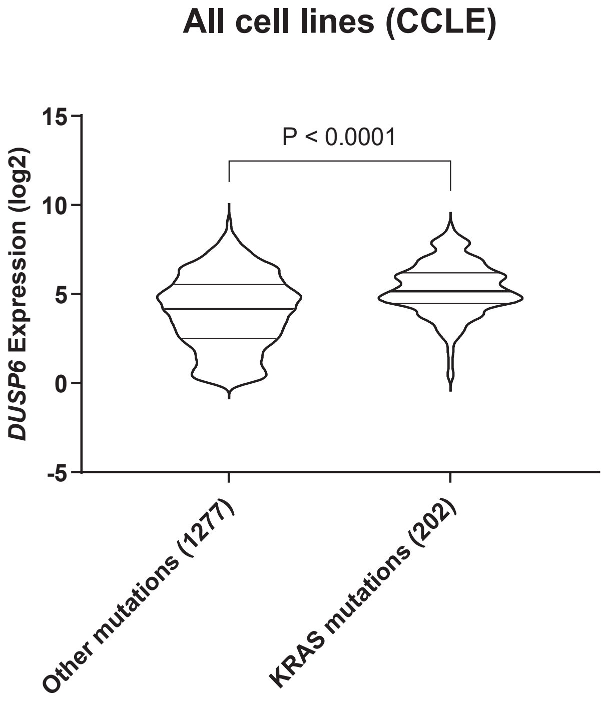

**Supplementary Figure 1 – A)** Pearson correlation analysis between *DUSP6* expression and *MAPK1* expression using the quasi-mesenchymal subset of GSE15471 dataset; **B)** *In silico* gene expression analysis utilizing GSE93326 to assess DUSP6 differential expression between the epithelial and the stromal compartment in primary tumor samples. Mann-Whitney unpaired test; **C)** Western blotting analysis assessing DUSP6 protein levels in human tumor cell lines, mouse tumor cell lines and cancer associated fibroblasts (CAF) cell lines; **D)** *Dusp6* expression in tumor sections derived from *LSL-Kras*$^{G12D/+}$, *Ptf1a*$^{Cre/+}$ mouse pancreatic tissue (KC) and *LSL-Kras*$^{G12D/+}$, Tp53$^{R172H/+}$, *Ptf1a*$^{Cre/+}$ mouse pancreatic tissue (KPC) using the RNAscope technology. *Dusp6* (green); *Krt19* (red, upper images); *Pdgfrb* (red, lower images); Pearson correlation analysis between *DUSP6* expression and *MAPK1* expression using the epithelial compartment **(E)** and the stromal compartment (F) of GSE93326 dataset.

**Supplementary Figure 2 – A)** *DUSP6* expression among cell types in healthy pancreatic tissue, adjacent normal pancreatic tissue and PDAC samples assessed by single-cell RNA-seq [28]. **B)** *DUSP6* expression was assessed by bulk RNA-seq (GSE245535) in liver biopsy samples from a cohort of patients who underwent pancreatectomy and were followed for 3 years to monitor metastasis development [40]. Mann-Whitney unpaired test.

**Supplementary Figure 3 –** Validation of DUSP6 knockdown in PDAC cell lines. DUSP6 levels were normalized with β-actin levels, while P-ERK1/2 levels were normalized with total ERK levels.

**Supplementary Figure 4 – DUSP6 downregulation reduces proliferation and migration in PDAC cell lines *in vitro*.** Assessment of proliferation and migratory capacity in AsPC-1 (**A** & **C**) and BxPC-3 (**B** & **D**) with DUSP6 knockdown. Graphs show 3 independent experiments with 4-6 replicates each. Results represent mean ± SEM; Brown-Forsythe and Welch ANOVA test. *ns*: not significant; ** P < 0.01; **** P < 0.0001.

**Supplementary Figure 5 – A & B)** Assessment of proliferation of DUSP6 knockdown cells upon 2-DG treatment. Seahorse Glycolysis Stress Test was performed in 9-12 replicates in 2 independent experiments; **C)** Extracellular acidification rate in PDAC cell lines upon combination of electron transport chain inhibition (Rotenone and Antimycin A) and glycolysis inhibition (2-DG). Media contains 1 mM pyruvate, 2 mM glutamine and 10 mM glucose; Results represent mean ± SD of one representative experiment. Welch's unpaired t-test;

Proliferation graphs show 3 independent experiments with 4-6 replicates each. Results represent mean ± SEM. Mixed-effects analysis with Šídák's multiple comparisons test. * $P < 0.05$; **** $P < 0.0001$.

**Supplementary Figure 6 –** *DUSP6* expression analysis performed *in silico* on the CCLE dataset. Data shows all the cell lines available in the platform, stratified into cells harboring KRAS mutations and cells that do not.
